# Downward Gazing for Steadiness

**DOI:** 10.1101/2020.02.28.969162

**Authors:** Yogev Koren, Rotem Mairon, Ilay Sofer, Yisrael Parmet, Ohad Ben-Shahar, Simona Bar-Haim

## Abstract

When walking on an uneven surface or complex terrain, humans tend to gaze downward. Previous investigations indicate that visual information can be used for online control of stepping. Behavioral investigations suggest that, during walking, the availability of visual information increases stepping accuracy, but probably through a feedforward control mechanism. Consequently, downward gazing (DWG) is usually interpreted as a strategy used to acquire useful information for online and/or feedforward control of stepping.

Visual information is not exclusively used for guiding locomotion; a wealth of literature has been published on the usefulness of visual information for feedback postural control. Critically, postural control has been shown to be sensitive to the visual flow arising from the respective motion of the individual and the 3D environment.

To investigate whether DWG can be used to enhance feedback control of posture, rather than feedforward/online control of gait, we conducted a series of experiments that explore this possible interplay. Through these experiments we were able to show that DWG, just a few steps ahead, results in a steadier standing and walking posture, without the need for accuracy. Moreover, we were able to demonstrate that humans resort to DWG when walking stability is compromised, even when destabilizing features were visually unpredictable.

This series of experiments provides sufficient evidence of the possible interplay between visual information used for guiding locomotion and that used for postural control. Moreover, this evidence raises concerns regarding the way we interpret gaze behavior without the knowledge of the type and use of the information gathered.

## Background

Visual information is important for guiding human locomotion (Patla, 1997; Marigold, 2008). When walking on uneven surface (Marigold and Patla, 2007) or complex terrain (Matthis, Yates and Hayhoe, 2018), healthy individuals tend to gaze downward, presumably to reduce the surface’s uncertainty (Domínguez-Zamora, Gunn and Marigold, 2018) by identifying individual footholds and use this information, in feedforward (Matthis, Barton and Fajen, 2015) and/or online (Reynolds and Day, 2005a; Reynolds and Day, 2005b; Smid and Den Otter, 2013), to control the leg’s trajectory. Given a well-established relation between gaze behavior and footholds in precise stepping paradigms (e.g. (Hollands *et al.*, 1995; Patla and Vickers, 2003)), and the increased stepping accuracy when foveal information (Smid and Den Otter, 2013) is available during swing phase (Reynolds and Day, 2005a) in such paradigms, this perspective is only logical. Evidence from similar paradigms, such as obstacle negotiation (Patla and Vickers, 1997; Matthis and Fajen, 2014) and stair climbing (Zietz and Hollands, 2009) extend this perspective to other situations where leg trajectory’s accuracy is important.

It has been suggested that the extent to which an individual fixates on a foothold depends on the perceived challenge to walking stability (Patla and Vickers, 2003); or more generally, fixation time depends on the perceived relevancy to the task (Domínguez-Zamora, Gunn and Marigold, 2018). Recent reports (Ellmers, Cocks and Young, 2019; Ellmers and Young, 2019) support such assumption, as they indicate that anxiety (fear of falling) also leads to downward gazing (DWG). This observation was interpreted, according to the above perspective, as an attempt for online, conscious, control. Taken together, these evidence suggest that DWG, when a person is walking on uneven or complex terrain, is an attempt to consciously control each individual step, triggered when walking stability is perceived as compromised.

Visual information is not used exclusively for guiding locomotion; it is also important for postural control (e.g. (Lee and Lishman, 1975; Stoffregen, 1985; Bardy, Warren and Kay, 1996; Guerraz *et al.*, 2000)). Previous reports suggest that both are greatly influenced by the visual structure of the environment (Warren, Kay and Yilmaz, 1996; Bardy, Warren and Kay, 1996; Warren *et al.*, 2001; Salinas, Wilken and Dingwell, 2017), specifically, the type, magnitude, and direction of sensory information arising from the visual flow. Not only ocular, but also extra- ocular information may be used for postural control. This information is thought to be most useful for short gaze distances (Guerraz and Bronstein, 2008), which is the case for DWG. Therefore, postural control may provide a complementary, and in some cases an alternative, explanation for DWG. In support of such a possibility, DWG has been reported to enhance postural steadiness for standing (Aoki *et al.*, 2014) and walking (Aoki *et al.*, 2017) stroke survivors.

To the best of our knowledge, excluding the above-mentioned report, no other attempts have been made to study DWG behavior in the context of postural control during walking. Moreover, the possible interplay between visual information needed for guiding locomotion and that needed for postural control is usually overlooked in behavioral studies. Thus, we wanted to test the hypothesis that DWG is not used exclusively for leg trajectory control, but may also serve as a way to alter the visual flow in order to gain useful information for feedback (online) postural control.

Walking stability is naturally an important outcome of such investigation. Since this term has no widely acceptable definition in the literature (Bruijn *et al.*, 2013), in this paper the term “stability” will be used under the definition, “the ability to perform without falling” (standing or walking) and the term “steadiness” under the definition “regularity and/or consistency of the motor output” (e.g. the magnitude of variance of a given parameter). No preliminary assumptions regarding the relation between these two terms have been made.

In this paper we present a series of experiments designed to explore the effect of DWG on the steadiness of healthy adults: first we tested how DWG affects standing postural steadiness, as reflected by the dynamics of center of mass (COM) motion. Next we tested how DWG effects the dynamics of the COM motion during walking, and finally we tested whether humans will resort to DWG even when the uncertainty of the walking surface cannot be visually resolved. The results of these experiments are then discussed in the context of the altered visual information during DWG.

Our hypotheses were that DWG will enhance postural steadiness of both standing and walking, and that our participants will resort to DWG despite the fact that DWG cannot resolve the surface’s uncertainty.

### Experiment 1

In this experiment we wanted to investigate the effect of DWG on standing postural steadiness. We started with a standing paradigm as postural steadiness is easily defined, as the motion of the center of mass (COM) about the fixed base of support (BOS), and has been shown to be sensitive to the visual flow arising from this motion. We recruited 15 healthy adults (7 males and 8 females, ranging from 20 to 41 years old, with a mean age of 28) and evaluated their postural sway under five visual conditions: eyes closed (EC), downward gazing at their feet (DWGF), downward gazing one meter ahead (DWG1), downward gazing three meters ahead (DWG3), and forward gazing (FG) at eye level approximately four meters ahead. Participants were instructed to stand as still as possible on a force-plate (Kistler 9286AA) in a standardized, narrow-base stance for 30 seconds. Each participant was tested five times under each condition in random order.

As outcome measures for postural steadiness, three parameters derived from the stabilogram diffusion analysis (SDA) were used. Specifically, the short-term displacement coefficient (originally the term ‘diffusion coefficient’ was used) of sway on the X-axis (D_xs_), on the Y-axis (D_ys_), and planar displacement (D_rs_) (Collins and De Luca, 1993). Although the mechanism underlying SDA is controversial (e.g. (Peterka, 2000)), it offers the possibility to investigate the time-dependent characteristics of postural sway. To test whether DWG alters the directionality of sway, we also calculated the difference between medio-lateral (ML) and anterior-posterior (AP) sway range (i.e. sway range difference, SRD). (For a full description see the *Methods* section). Results of this experiment demonstrated a significant main effect (F= 65.8-91.2, p<0.001) for the visual condition in all SDA parameters. As expected, when visual information was unavailable (i.e. EC) sway values of all parameters were the greatest, signifying that visual information is important for postural control (see Figure 1). When visual information was available (i.e. excluding EC condition), minimal mean sway values, of all parameters, were observed during the DWG1 and DWG3 conditions (for pairwise comparison see Figure 1). Results of the directionality testing revealed a main effect for the condition (F=5.393, p<0.001). Individual results revealed that mean SRD values in all DWG conditions were not different from zero, as indicated by the 95% CI of the mean (DWGF -4.15-1.6, DWG1 -0.81-4.95, DWG3 -0.91-4.85), suggesting isotropic sway. For both EC and FG conditions, mean values were significantly greater than zero, as indicated by the 95% CI of the mean (EC 2.51-8.27, FG 0.63-6.39), indicating sway on the ML axis was greater than on the AP axis.

**Figure 1.**
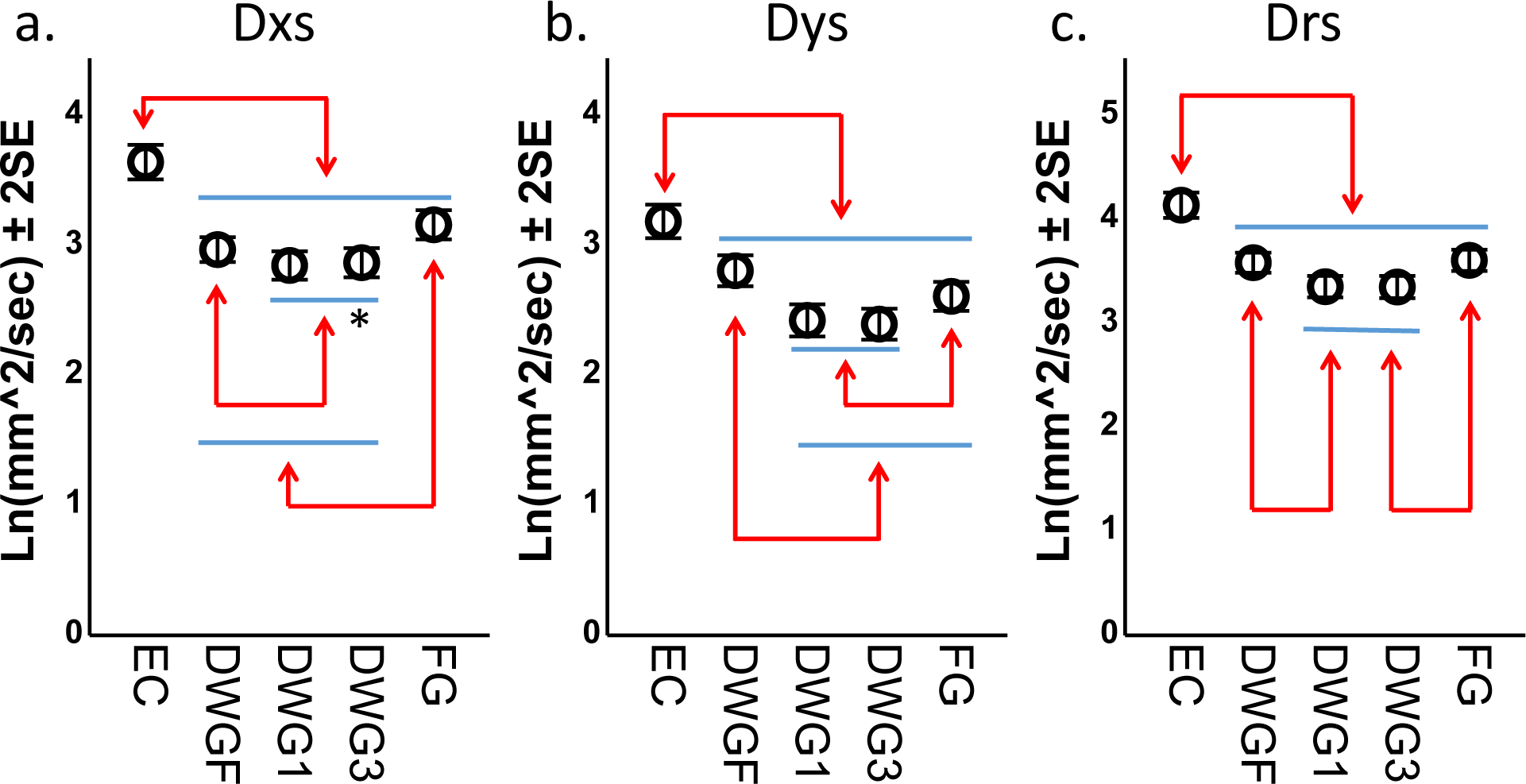
pairwise comparison of mean short-term displacement coefficient values (log-transformed) by condition for the parameters *D_xs_ (a), D_ys_* (b) and *D_rs_* (c). Arrows indicate significant differences at the level of α<0.05. * indicate p=0.055.

The main purpose of this experiment was to investigate the effect of downward gazing on standing postural steadiness. Our results clearly demonstrate that posture is affected by visual information as evident from the consistent highest values achieved during the EC condition. Excluding the EC condition, gaze position also had a significant effect on postural sway. Generally, DWG1 and DWG3 yielded equivalent results for all parameters. Mean values during these conditions were consistently the lowest, indicating greatest postural steadiness. These results, therefore, support our hypothesis that DWG may be used to enhance postural steadiness. Moreover, the directionality results support our assumption that this effect was achieved due to the altered visual structure and the associated change to the visual flow (see *Discussion* section).

### Experiment 2

Next we wanted to assess whether DWG will also affect walking postural steadiness, which was our main research question. To answer this question we tested 30 participants (15 males and 15 females, ranging from 20 to 45 years old, with a mean age of 27.6) under 5 visual walking conditions, four of which were similar to those in the standing postural testing (i.e. DWGF, DWG1, DWG3, FG). The fifth condition, which served as baseline in this experiment, was unrestricted (UR) in which participants were free to act as they please. For DWGF, participants were instructed to look where they assumed their next step is going to be, which for treadmill walking is about half a step forward. All tests were performed using a treadmill with an embedded force-plate (ForceLink, Culemborg, The Netherlands) able to record vertical ground reaction force (GRF) and the trajectory of the center of pressure (COP). Participants were instructed to walk, at their preferred velocity, for four minutes in each condition. Tests were executed continuously with a 10-sec break between conditions, during which participants were instructed of their next gaze position, the order of which was random. The only instructions given were “look continuously on the specific target of the condition,” and “try to maintain your position in the middle of the treadmill but without looking.”

As outcome measures we quantified the steadiness of the COM motion as estimated from the GRF time series; specifically, we computed the local divergence exponent (λ*) for the vertical GRF (Fz), COPX and COPY time series. The divergence exponent is a commonly used parameter thought to reflect the local steadiness of walking (Dingwell and Cusumano, 2000). Like SDA, λ* also captures the time-dependent characteristics of the motor output. As secondary outcome measures, we calculated the variability (median absolute deviation, MAD) of the parameters step-width and step-time.

First we evaluated whether participants maintained their intended, anterior-posterior (AP), position on the treadmill (which was determined beforehand by placing a 5 kg weight in the middle of the treadmill and recording its position in the force plate coordinates) during the walks and whether velocities were different between conditions (see Figure 2). Results of the position testing revealed a main effect for the condition (F=6.3, p<0.001). Specifically, values during DWGF and DWG1 were greater than values during baseline condition (UR) with a maximal mean difference of 7 cm. However, only in the DWG1 condition did mean value significantly differ from the intended position (mean=5.9 cm, 95%CI 2.4-9.4 cm). Results of the velocity testing also revealed a main effect for the condition (F=2.6, p=0.038). Specifically, mean velocity during the DWGF condition was slower than velocities in the other conditions with a mean difference of 0.09 m/s from baseline (UR).

**Figure 2:**
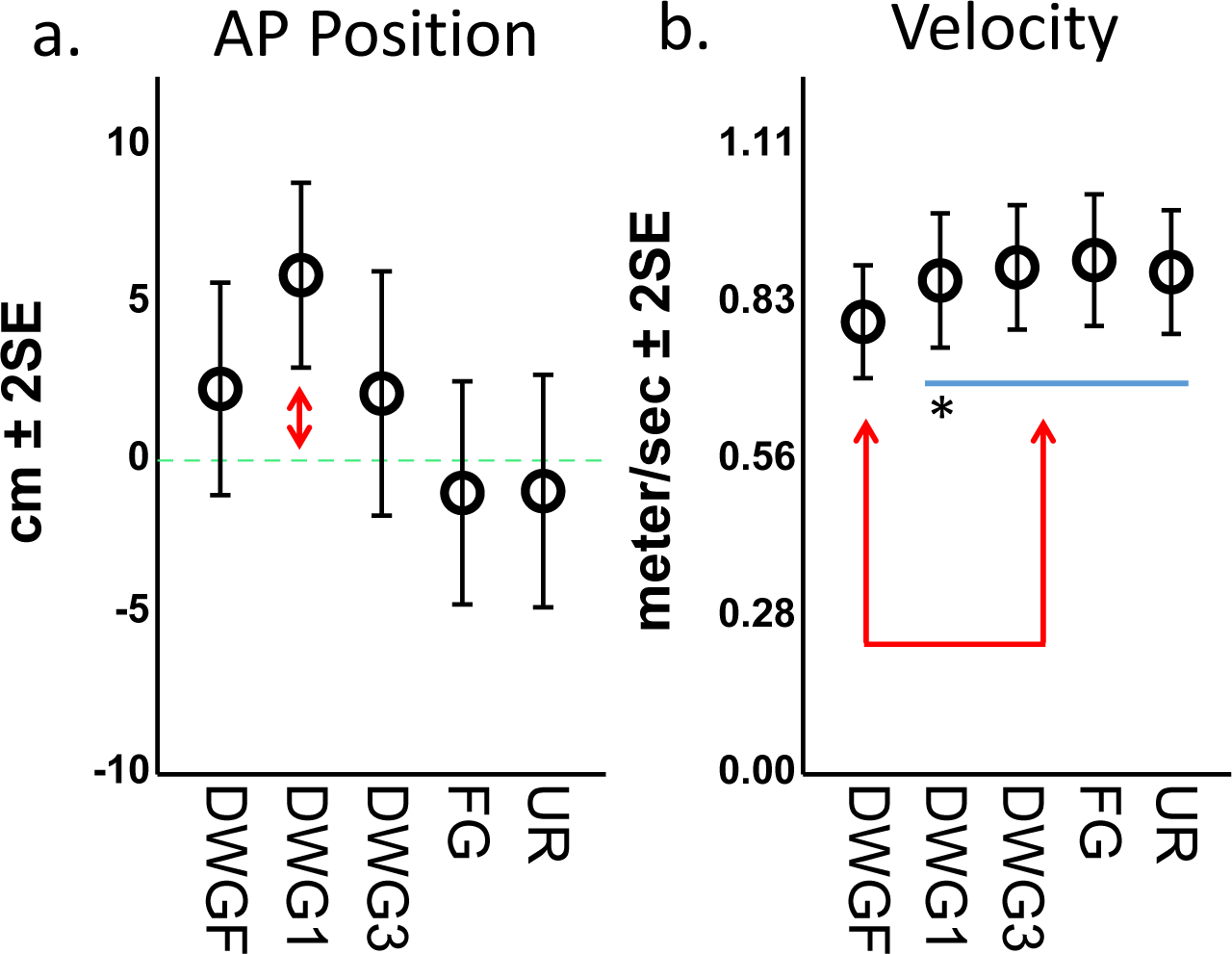
mean AP center of pressure position (a) relative to the intended position by condition, and pairwise comparison of the preferred walking velocities (b) by condition. Arrows indicate a significant difference at the level of α<0.05. * indicate p=0.054.

As for our main outcome measures, ML and AP COM motion steadiness, as indicated by the parameters λ*COPX and λ*COPY, was unaffected by gaze position (F=1.5, p=0.2 and F=2.0, p=0.1 respectively), but steadiness of the vertical component, as indicated by the parameter λ*Fz, was significantly affected by gaze position (F=5.4, p<0.001). To estimate whether λ* values were affected by walking velocity, as was previously reported (Dingwell and Cusumano, 2000; England and Granata, 2007), we tested these value against the self-selected velocity and found significant correlations with λ*COPY and λ*Fz values (F=12.0, p=0.001 and F=51.1, p<0.001 respectively). Therefore, we corrected for velocity by including this parameter as a *fixed* factor within the λ*COPY and λ*Fz statistical models. Results of the velocity controlled models revealed that both parameters were significantly affected by gaze position (F=2.6, p=0.04; F=4.2, p=0.003 respectively). In both models, minimal values were observed during the DWG3 condition, but these were not significantly different from values during the DWG1 condition (for complete pairwise comparison see Figure 3). Values of the parameter λ*COPX were not correlated with velocity, and this parameter was not further explored.

**Figure 3:**
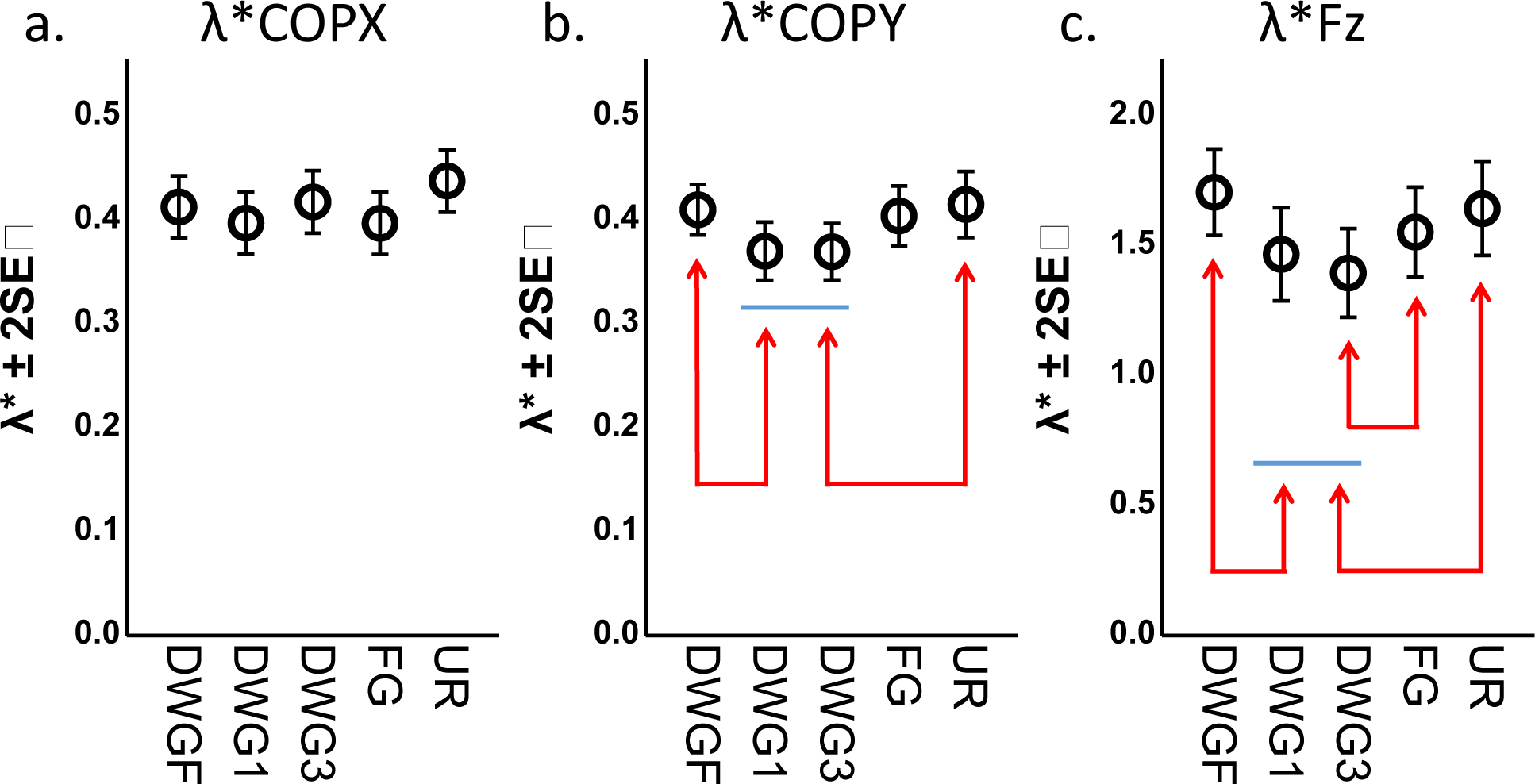
pairwise comparison of mean divergence rate (λ*) by gaze position of the *COPX (a), COPY (b)* and *Fz (c)* time series. For the latter two, velocity controlled predicted values are presented. Arrows indicate significant differences at the level of α<0.05.

To estimate walking steadiness we also used more traditional parameters, i.e. step-time and step-width variability (as the median absolute deviation, MAD). Results of these models did not reveal a significant main effect for the condition (F=0.6, p=0.66; F=0.5, p=0.74 respectively). The former was significantly affected by walking velocity (F=131.1, p<0.001), and the velocity-controlled model revealed minimal mean value during the DWG3 condition. However, this effect did not reach significance level (F=1.7, p=0.15).

These results partially support our hypothesis, as DWG resulted in a steadier motion of the COM in the AP, but not the ML, direction. Moreover, results of the parameter λ*Fz also matched our predictions. Since the time-dependent change in ground reaction force (i.e. Fz) primarily reflects the acceleration of the COM on the vertical axis during walking, these results suggest that DWG enhances postural steadiness also along the vertical axis. We believe that this is achieved due to visual expansion and motion parallax created by the vertical motion of the COM (see *Discussion* section). However, this steadiness was not translated into a steadier gait, as reflected by step variability parameters.

### Experiment 3

As was mentioned earlier, DWG is usually interpreted as a strategy used to gather information to control each individual step when stability is perceived as compromised. Above we have provided evidence that DWG enhances postural steadiness of both standing and walking. This evidence suggests an alternative (or maybe complementary) explanation for DWG. However, since we made no assumptions regarding the relation between steadiness and stability, such an alternative is reasonable only if steadiness is beneficial or otherwise desirable. Therefore, we wanted to investigate whether steadiness is desired when stability is compromised. In other words, do humans resort to DWG even when visual information cannot resolve the uncertainty problem?

To answer this question we used an experimental paradigm in which participants walked with and without perturbations while their gaze behavior was monitored. To ensure participants are unable to use vision to resolve the perturbation problem, these were induced using the Re-Step system (Bar-Haim *et al.*, 2013; Koren *et al.*, 2018) (see Figure 4). Briefly, the Re-Step (RS) is a shoe-like system able to continuously change, during the swing phase, the shape of its sole. These changes are perceived, during the stance phase, as walking on uneven surface. As these changes take place underneath the sole, they are visually unpredictable. Therefore, under the common interpretation, DWG is not expected when walking with the RS system.

**Figure 4:**
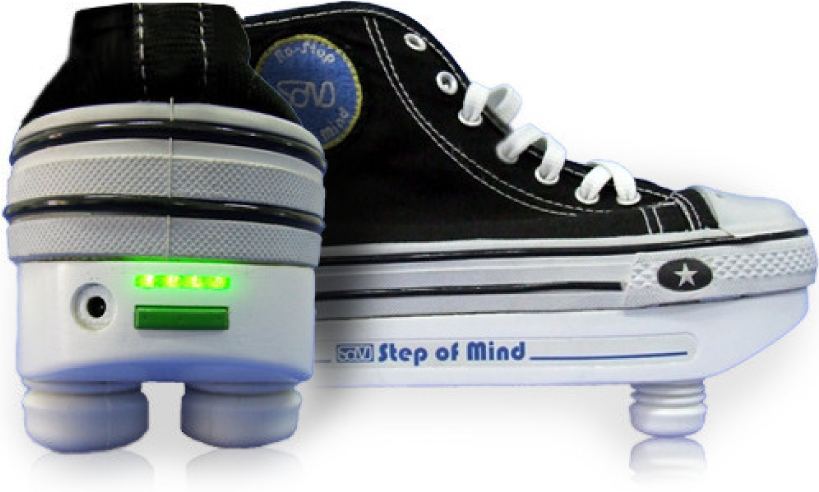
the Re-Step is a shoe-like system able to change, continuously, the shape of its sole. The pistons, underneath the sole, change their length during the swing phase, creating a plane oblique to the sole’s plane. These changes are visually unpredictable and perceived as walking on uneven surface.

**Figure 5:**
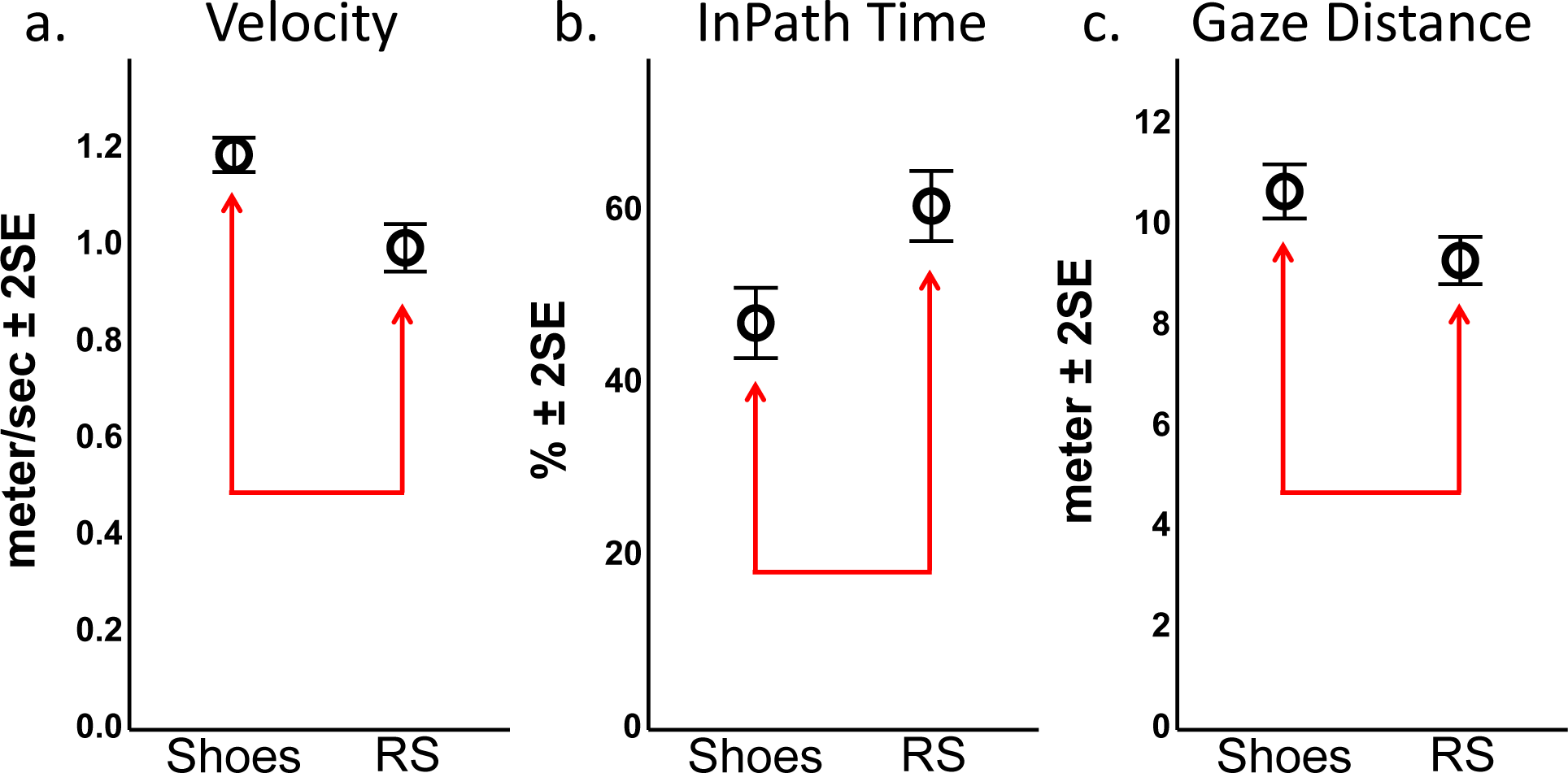
comparison between shoe types of (a) walking velocity, (b) percentage of time spent looking onto the future path (i.e. InPath), and (c) gaze distance. Arrows indicate a significant difference at the level of α<0.05.

We recruited 15 healthy adults (mean age 25, range 20-30) and monitored their gaze position under two walking conditions: walking with their own shoes and walking with the RS system. Participants performed six 20-meter long walks in each shoe type. All tests were performed in the same, well lit, corridor. Gaze position was continuously monitored using binocular eye-tracking glasses (SensoMotoric Instruments, Teltow, Germany) at a 60Hz sampling rate. Gaze data were referenced to the vanishing point, which was computed using a dedicated algorithm and software. Further, based on these computations we were able to calculate gaze distance whenever fixations were made onto the walking path (i.e. the real or imaginary future walking surface). We used gaze position distance from the vanishing point, on the vertical axis (vertical gaze position), indicating DWG extent, as our main outcome measure. For simplicity, we used the inverse of these values such that large values indicate a gaze directed straight ahead (i.e. toward the horizon) and small values indicate DWG. Secondary outcome measures included the look-ahead distance and its variance. Walking velocity and the proportion of gaze positions (from the total number of samples) directed onto the future path were also calculated. For a more detailed investigation, we divided gaze distances into three categories: up to 3 meters ahead (true DWG), 3-10 meters ahead, and more than 10 meters ahead. We then tested the effect of the shoe type on the proportion of time spent looking within each category. (For a complete description, see the *Methods* section).

We hypothesized that walking with the RS system will lead to a more cautious gait (as previously observed (Koren, Parmet and Bar-Haim, 2019)), as indicated by a decrease in walking velocity. This change will be accompanied by an increase in DWG, i.e. an increase in the vertical distance from the vanishing point (as indicated by smaller values) and a decrease in gaze distance.

Results of this experiment demonstrated a significant main effect for the shoe type in all outcome measures: walking velocity significantly decreased in the RS condition (MD=-0.19 ± 0.01 SE, p<0.001). Proportion of time spent looking onto the walking path significantly increased (MD=13.6 ± 2.1 SE, p<0.001). Look ahead distance (MD=-1.3 ± 0.43 SE, p=0.004) and its variance (MD=-0.97 ± 0.26 SE, p<0.001) significantly decreased during walks made with the RS.

As for our main outcome measure (see Figure 6), a significant increase in DWG extent (MD=8.2 ± 1.2 SE, p<0.001) was observed during walks with the RS. When we tested the effect of the RS system in each of the distance categories (see Figure 6), a significant increase (MD=11.6 ± 1.9 SE, p<0.001) was observed in the mid-range (i.e. 3-10 meters ahead) category, and no change in the short- and long-range (i.e. up to 3 and more than 10 meters) categories (MD=1.3 ± 1.9 SE, P=0.5; MD=0.77 ± 1.9, P=0.69 respectively).

**Figure 6:**
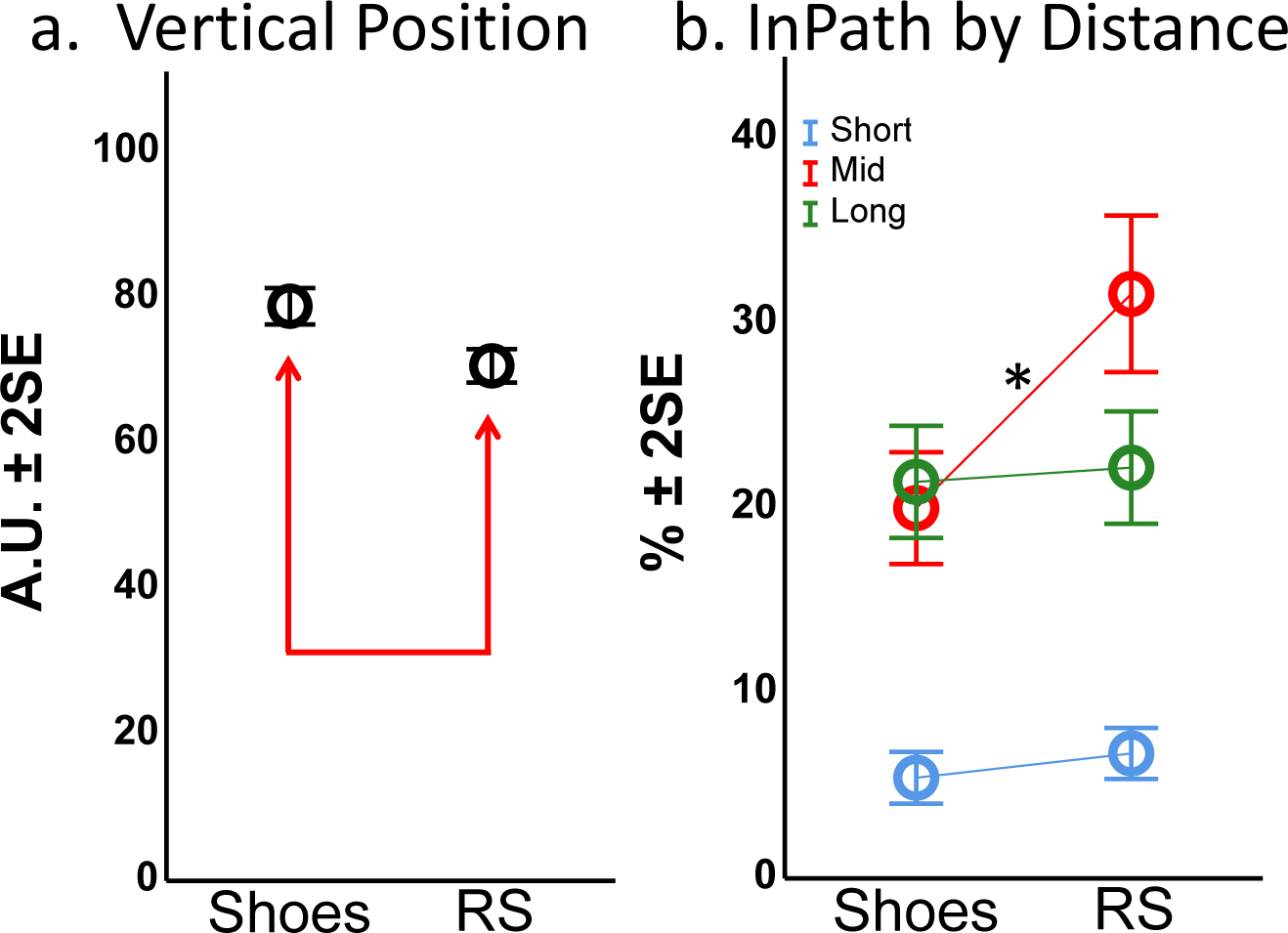
pairwise comparison of the vertical gaze position (a) and the percentage of time spent looking onto the future path (i.e. InPath) within each distance category (b) by shoe type. Arrows and * indicate a significant difference at the level of α<0.05.

These results support our hypothesis, as walking velocity decreased, suggesting walking stability was perceived as compromised, vertical gaze distance decreased and look ahead distance decreased, as expected. However, the increase in time spent looking onto the future path was primarily due to change in gaze behavior in the mid-range category, i.e. distances between 3-10 meters ahead, and not into “true” DWG, up to 3 meters ahead, as expected.

## Discussion

The main purpose of this paper was to investigate the effect of DWG on walking steadiness. During walking, DWG is usually interpreted as an attempt at online or feedforward control of stepping. By contrast (or maybe in conjunction), DWG has been shown to affect standing postural steadiness. Clark (1995) reported that when toddlers were provided with external trunk support an adult-like walking pattern emerged. This was interpreted, from a dynamical systems perspective, as an indication that postural control is an important constraint on behavior. Indeed, Kay and Warren (2001) reported a complex relation between gait and posture that was sensitive to visual perturbations. Therefore, when considering how visual information is used to guide locomotion, one also has to consider the interplay with postural control.

Previous mechanistic investigations suggest that the combined biomechanical and vestibular effect of downward head inclination (i.e. neck flexion) decreases postural steadiness (Buckley *et al.*, 2005), but downward eye movement (i.e. eyes in head) was reported to increase postural steadiness (Kapoula and Le, 2006; Ustinova and Perkins, 2011). These seemingly contradicting reports suggest that the need to control individual steps may come with the cost (or benefit) of reduced postural steadiness or with the benefit (or cost) of enhanced postural steadiness. In the first experiment, our results for the parameter Dxs indicate that DWG reduced ML sway in comparison to the FG condition. On the other hand, AP sway, as indicated by the parameter Dys, increased during the DWGF condition but decreased during DWG1 and DWG3 conditions in comparison with the FG condition. The overall planar sway, as indicated by the parameter Drs, was reduced during the DWG1 and DWG3 conditions in comparison to the FG condition. However, planar sway during the DWGF condition did not differ from the FG condition.

From a visual information perspective these results may be explained by the change in visual flow induced by the different gaze positions. For the purpose of this discussion we will consider three variables of the optical flow generated by the motion of the self relative to the environment: simple flow, which refers to planar or lamellar flow (parallel to the observer); motion parallax, which refers to differential flow due to parallax (most prominent when gaze is orthogonal to the motion direction); and radial expansion, which refers to a flow due to a differential rate of expansion (most prominent when gaze is in the direction of motion). For the latter, the term visual expansion seems more appropriate to the current discussion, since in most cases we consider the increase/decrease in magnitude of a single, two-dimensional, object as its distance from the observer changes. Such scale changes seem sufficient to perceive motion (Schrater, Knill and Simoncelli, 2001). It has been suggested that postural control is more sensitive to the last two variables (Warren, Kay and Yilmaz, 1996; Bardy, Warren and Kay, 1996; Stoffregen, 1985; Guerraz *et al.*, 2000) and that this sensitivity increases with flow density (Warren Jr, 1995). Therefore, this information will be considered superior to simple flow, and gaze directed at the support surface (i.e. DWG) is assumed to contain more visual cues (due to tiles, markings, patterns, etc.).

In the DWG1 and DWG3 conditions, gaze is diagonal to the targets and therefore both near and distant features of the surface are contained within the visual field (around the target). This visual structure includes depth and will generate motion parallax for ML sway and visual expansion for AP sway (Warren, Kay and Yilmaz, 1996; Guerraz *et al.*, 2000). Thus, in both conditions the visual flow generated by sway in any given direction is adequate for feedback control, and therefore equivocal and isotropic sway is expected, as was observed. During the DWGF condition, gaze is perpendicular to the viewing target and provides a planar visual structure. Sway in this visual structure will generate simple flow in any given direction, which is less adequate for postural control than motion parallax and visual expansion. Therefore, sway in this condition is expected to be isotropic and increase in comparison to the DWG1 and DWG3 conditions, as was observed. FG condition, in the current study, also provides a planar visual structure as the target was placed on the back wall of the laboratory. Moreover, this was a white wall offering no or a minimal amount of visual cues. Sway in this condition will generate visual expansion in the AP direction and simple flow in the ML direction. Therefore, sway is expected to be anisotropic, with reduced sway in the AP direction, as was observed. AP sway in this condition is also expected to be reduced compared to the DWGF condition, which was observed, but equivocal with sway in the DWG1 and DWG3 conditions, which was not observed. ML sway in this condition is expected to be greater than in the DWG1 and DWG3 conditions, as was observed, but equivocal with sway in the DWGF condition, an outcome that was not observed. Both inconsistencies can be explained by the lower visual cue density, provided by the wall.

Not only ocular, but also extra-ocular information affects postural sway (Guerraz and Bronstein, 2008). This information is sensitive to gaze distance (Guerraz *et al.*, 2000; Kapoula and Le, 2006) and may provide an alternative explanation of our results. However, these previous mechanistic investigations had used gaze distances irrelevant to walking. Moreover, under normal sway values, this information quickly decays to sub-threshold levels (Guerraz and Bronstein, 2008). A comparison of DWG1 and DWG3 conditions, in which the visual structure was similar but gaze distance differed, supports this notion, as for all parameters tested no difference was found between them. However, in our DWGF condition, gaze distance was short enough to provide extra-ocular information and therefore may provide an additional, or alternative, explanation to the reduced ML sway in this condition compared with FG.

Contrary to our results, Aoki et al. (2014) observed an increase in ML sway, but not for AP, when healthy elderly gazed downward. After controlling for gaze distance, both AP and ML sway increased in comparison to FG. By contrast, DWG decreased AP sway (with and without distance control) and ML sway (only without distance control) in stroke survivors. Although the inconsistency with our results may be related to methodological differences, it is also quite possible that age and pathology alter the effect of DWG. Specifically, Aoki et al. suggested reweighing of sensory information to explain their results. As mentioned earlier, Buckley (2005) also reported increased sway during DWG. However, Buckley intentionally maintained the visual structure of the environment, to control for differential flow. We therefore interpret their results not as contrary to our own but as supportive of our hypothesis that DWG can be used to alter the visual structure of the environment, and the associated visual flow, to enhance postural control.

That being said, in this experiment we did not try to investigate a specific mechanism(s), but to establish the overall effect of DWG to commonly reported distances (Patla and Vickers, 2003; Marigold and Patla, 2007; Matthis, Yates and Hayhoe, 2018) during walking. Results of this experiment were unambiguous, demonstrating that DWG to one and three meters ahead (roughly corresponding to 1-1.5 and 3-4 steps ahead, respectively) significantly increased postural steadiness. Although we suggest this effect was primarily derived from changes in visual information, other mechanisms/influences, such as biomechanical, proprioceptive and vestibular, are quite possible.

For walking, we were able to identify only a single investigation of DWG’s effect on postural steadiness (Aoki *et al.*, 2017). According to this report, acceleration of the lumbar spine (at the level of L3 vertebra) was reduced in all three dimensions (i.e. vertical, medio-lateral, and anterior-posterior) when healthy older adults gazed down as opposed to forward, suggesting that DWG increases postural steadiness not only for standing but also for walking. Our second experiment was designed to explore this possibility.

As in the first experiment, the visual flow, created by self-motion, was expected to differ among conditions. Specifically, we assumed that the visual flow would be similar to that created by standing sway on the horizontal plane (i.e. AP and ML). In addition, the visual flow created by vertical motion is also expected to differ among conditions. Vertical motion can create both motion parallax and visual expansion. The former is most prominent when gaze is parallel, and the latter when gaze is perpendicular, to the walking surface (assuming visual cues are within the visual field); thus DWG just a few steps ahead (i.e. DWG1 and DWG3), as opposed to the DWGF and FG conditions, provides both motion parallax- and visual expansion-related information.

The results of the second experiment were complex and not as straight forward as those of our standing experiment. First, we found that the preferred walking velocity in the DWGF condition was slower than in the other conditions (except for DWG1 in which a trend was noted, p=0.054). We believe this was due to our instructions, which made participants attentive to each step (i.e. conscious control), thus, forcing additional task-related constraints. Next we noted that participants increased gaze distance in the DWG1 condition by drifting backward on the treadmill, suggesting they were uncomfortable with the constraints of this condition. Gaze distance alone cannot explain this observation as gaze distance in the DWGF condition was shorter. We believe participants were trying (unconsciously) to optimize a certain parameter/s by increasing gaze distance. Whether this parameter is related to optical flow or not is unclear at this time.

The main results of this experiment indicate that gaze position affected the steadiness of neither the ML nor AP motion, as indicated by the parameters λ*COPX and λ*COPY respectively. Further, we also used spatio-temporal parameters, which are more commonly used, and found that neither step-time variability nor step-width variability were affected by gaze position. Thus, overall, and as opposed to standing, walking steadiness on the horizontal plane was unaffected by gaze position. That said, the walking task goals did not require steadiness beyond what was perceived by participants as essential. In the standing task, participants were specifically instructed to stand as still as possible, making steadiness a task goal. In a recent review (Hayhoe and Matthis, 2018), the authors argued that walking gaze behavior is goal-directed and that its role is to provide relevant information to satisfy the task’s goal/s. Therefore, it is only reasonable that even if the visual flow contained information to enhance steadiness, task goals did not require or encourage participants to use this information. Moreover, after controlling for walking velocity, we found that AP steadiness was significantly affected by gaze position. Results of this model indicated that gazing downward to 1 and 3 meters ahead increased steadiness of the AP motion, thus suggesting that participants chose a walking velocity that enabled them to maintain AP motion steadiness under the constraints of the different conditions.

In addition to AP and ML motion, walking includes a vertical component. To estimate the steadiness of the latter we also calculated λ* for the ground-reaction-force time series (i.e. λ*Fz). Although this time series is not a trajectory per-se, it reflects the vertical acceleration of the COM associated with gait cycle. Since acceleration is the 2^nd^ derivative of position, we consider this time series as a proxy to position. Results of the λ*Fz statistical model indicate that gaze position affected the steadiness of the vertical motion. Specifically, we found that DWG to 3 meters ahead resulted in minimal λ*Fz values, indicating maximal steadiness. Controlling for velocity within the λ*Fz model resulted in minimal, and equivocal, values during the DWG1 and DWG3 conditions, as was observed in the λ*COPY model. Gazing downward just two steps (one stride) ahead was suggested (Matthis and Fajen, 2013) to be sufficient for feedforward control of the inverted-pendulum-like motion of the COM (this arc-like motion includes both vertical and AP components). These authors argue that such feedforward control enables precise stepping while exploiting the biomechanics of bipedal gait for energetic efficiency. However, this inverted pendulum model assumes a fixed distance of the COM from the BOS, which is not the case for humans due to knee joint motion. Thus, the vertical non-pendulum-dependent motion (i.e. knee flexion/extension), and its associated pendulum-dependent AP motion (i.e. change in the magnitude of forward motion due to knee flexion/extension) must be controlled throughout the single support phase to ensure the precision of the swinging leg. Our results suggest that such feedback control may be achieved by exploiting the visual flow created during DWG. Moreover, in the present experiment step precision was not required, but steadiness of the COM nevertheless increased when participants gazed downward just a few steps ahead, suggesting that DWG may be used for other purposes regardless of whether step trajectory is important. The idea itself is not new, and several authors have suggested that travel-gaze, (i.e. a prominent gaze behavior in which gaze travels at the walking velocity a constant distance ahead) may be used for postural control (Patla and Vickers, 2003; Zietz and Hollands, 2009). Since our treadmill paradigm more closely resembles travel-gaze than other gaze behaviors, we believe our results support these previous suggestions.

One interesting, or maybe puzzling, finding from our treadmill experiment was that steadiness during the UR condition was reduced (i.e. elevated values) in comparison to other conditions (i.e. DWG1 and DWG3). This finding suggests that steadiness might be unnecessary since participants were able to but chose not to use the strategy that maximizes steadiness.

Overall, the findings from both our standing and treadmill-walking experiments indicate that DWG increases postural steadiness, but that steadiness is not necessarily beneficial or otherwise desirable. Therefore, the common interpretation of DWG behavior (i.e. conscious control over leg’s trajectory) seems more likely than postural control. However, for healthy adults treadmill-walking does not pose a considerable challenge, specifically, a challenge to stability which is associated with DWG. Our third experiment (i.e. free-walking) was conducted to examine whether steadiness is desirable when stability is compromised (and indirectly establishing whether stability and steadiness are related). Results of this experiment showed that when walking was perturbed, participants looked a shorter distance ahead. Moreover, the vertical distance of gaze position from the vanishing point was greater (as indicated by smaller values) when walking with the RS system. Although these measures seem equivalent, they are not. Specifically, gaze distance was calculated only if the gaze was directed onto the future path. The vertical distance was calculated for all data points. The time participants spent looking outside the boundaries of the future path was by no means negligible (53% and 40% for the shoes and RS conditions respectively). Therefore, although DWG onto the future path can be interpreted as habitual behavior, DWG to areas on which the participant is not going to step cannot be interpreted as such. Previous observations (Wolsley, Sakellari and Bronstein, 1996) indicate that the perception of self-motion, along with the associated postural response, is not “hard-wired” with a specific variable of the optic flow, but is flexible and is modulated by information about eye-in-head and head-on-body position. Thus, if DWG is used for postural control, where participants looked (i.e. forward or sideways) should not have changed their ability to use the visual flow created by DWG for postural control. We therefore interpret these results as an indication that DWG, and therefore postural steadiness, is desirable (when stability is compromised) whether or not it provides visual information about future footholds or obstacles.

Two more interesting findings arise from our free-walking experiment: 1) gaze distance variability decreased in the RS condition, and 2) the greatest effect was observed in the 3-10- meter gaze distance category and not as expected (i.e. up-to-3-meter category). We believe this is not just a reflection of the statistical relation between the two, but an indication that a new gaze strategy was adopted. Specifically, participants adapted by using a travel-gaze strategy (as indicated by the decrease in variability) to a distance that enabled them to increase postural steadiness while satisfying other task goals. Although this distance was not within the range that we expected, there is no reason why DWG to other distances should not provide the same visual information (depending on internal and external factors such as visual equity and saliency of visual cues). We do acknowledge that this last statement (or interpretation) does not necessarily reflect the true nature of our observation and should be more carefully examined. Moreover, this experiment is a behavioral investigation and our mechanistic interpretation is merely a suggestion. A future mechanistic investigation is guaranteed.

### Summary

In this series of experiments we were able to demonstrate that postural steadiness (as indicated by the dynamics of the COM) can be enhanced by gazing down just a few steps ahead. However, steadiness is not necessarily desirable, unless otherwise dictated by a task’s constraints or goals. Stability might be one such constraint, as DWG was observed when it was compromised even though the uncertainty of the walking surface could not be visually resolved. The series of experiments presented here is not intended to challenge current perspectives in any way. We simply wish to argue that visual information, even if directed at the walking surface, may serve other purposes, such as controlling posture and should be interpreted carefully.

## Methods

Participants in all experiments were healthy adults, recruited primarily from the university’s undergraduate programs. Advertisement was primarily through word of mouth and social media. In all cases participants were paid 11-17 USD, depending on the experiment’s duration, in local currency. All experiments were approved, beforehand, by the appropriate ethics committee. Participants were informed about purpose and methods beforehand but information regarding the hypothesis was intentionally omitted. After they agreed to participate, written informed consent was obtained and demographic data collected. For descriptive statistics of these data, see Table 1 below.

In all cases, statistical analysis was performed using linear mixed-effect models with participants as the *random* effect. Significance level was set to α=0.05, and the least significant difference (LSD) method was used to correct for multiple comparisons.

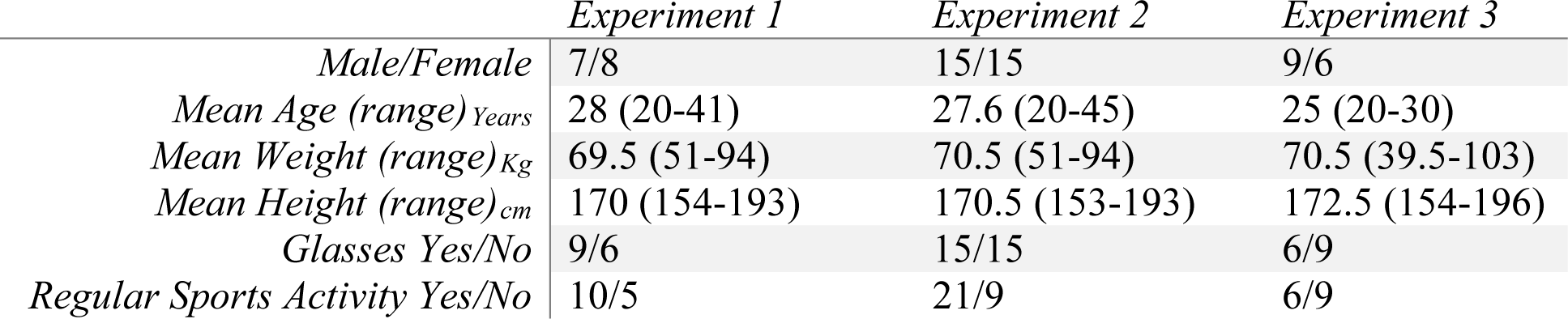

*Experiment 1-* Posturography of fifteen adults, reporting no neurological or other condition affecting gait, was evaluated in this experiment. Participants were tested under five visual conditions: eyes closed (EC), downward gazing at their own feet/toes (DWGF), downward gazing one meter ahead (DWG1), downward gazing three meters ahead (DWG3) and forward gazing (FG) at a target approximately 4.2 meters ahead at eye level.

Participants were instructed to stand barefoot, as still as possible, on a Kistler 9286AA force platform (Kistler Instrument Corp., Winterthur, Switzerland) in a standardized stance, i.e. with their feet tight together and hands loosely hanging at their sides. Five 30-second quiet-standing trials in each of the five gaze positions were performed (with a total of 25 trials in random order). Raw data from the force plate were collected, at 100Hz, using a data acquisition system consisting of a data acquisition box (Kistler A/D type 5691) and Bioware (version 5.3.0.7) software.

Before each trial (i.e. a single 30-sec stand) the force plate was calibrated with no weight (i.e. participants were instructed to step off the platform). Following the calibration, participants were instructed to stand on the platform and continuously look at one of the five targets. Locations for DWG1, DWG3 and FG were marked with colored circles 20 cm in diameter. For the DWGF, participants were instructed to look at their own toes, while for the EC condition, no specific instructions were given besides “close your eyes”.

The recordings were then processed by a dedicated MATLAB script. First, the center of pressure (COP) time series was low-passed using a 2nd order Butterworth filter with a cutoff frequency of 15Hz. The script computes the short-term displacement coefficient (D_is_) of COP driven from stabilogram diffusion analysis (SDA) as described by Collins and De-Luca (1993). Briefly, the displacement (originally the term “diffusion” was used) coefficient is the rate at which the quadratic Euclidean distance between two COP positions increases as a function of the time interval between them. That is, for a given Δt, spanning m data intervals and N samples, planar displacement (Δr^2^) is calculated as:

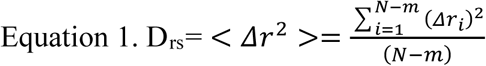

This calculation is repeated for every Δt, and the D_is_ is calculated as the slope of the Δi^2^ by Δt plot. In this experiment we calculated three coefficients: single dimension on the X (D_xs_) and Y (D_ys_) axes, and the planar coefficient (D_rs_), all given in mm^2^/sec.

In general, smaller sway values indicate increased steadiness and are usually considered an indication of better postural control, while larger values are thought to be indicative of decreased steadiness due to impaired control. This perspective is derived from empirical evidence showing increase in sway when balance control is assumed to be reduced, such as when comparing sway with or without visual information (Prieto *et al.*, 1996), young and older adults (Collins *et al.*, 1995), or elderly non-fallers and fallers (Melzer, Benjuya and Kaplanski, 2004; Melzer, Kurz and Oddsson, 2010).

To test isotropy, the script also computed sway range in the AP (COPY) and ML (COPX) directions (i.e. the distance between the extreme values in each direction) of each trial. The parameter Sway Range Difference (SRD) was then calculated as COPX-COPY.

For statistical analysis we used all available measurements (i.e. every trial was represented as a data point). A total of 375 trials were obtained, two of which were excluded; in the first case the participant scratched his head during the first DWG1 and in the second case (a different participant, at the third FG) the pre-trial calibration was performed incorrectly resulting in unreasonable outcomes (e.g. sway velocity of 410 m/s). These excluded values were replaced with the mean values of the participant in the specific condition. Since sway parameters’ distribution significantly deviated from normal distribution (excluding SRD), we used a logarithmic transformation (denoted as Ln(*parameter*)). The transformed values were analyzed using a linear mixed-effect model, with condition as the *fixed* effect and subject as the *random* effect. Residuals of the models were evaluated for normal distribution.

*Experiment 2-* thirty healthy adults were tested under five visual conditions: DWGF, DWG1, DWG3, FG and unrestricted (UR). For DWGF, participants were instructed to look where they assumed their next step is going to be, which for treadmill walking is about half a step forward. Participants were instructed to walk on a treadmill for 4 minutes, at their preferred walking velocity, while fixating on one of four visual cues (i.e. DWGF, DWG1, DWG3, FG) or simply walking without any instructions (i.e. UR). Before testing, participants selected their preferred walking velocity in each of the conditions. For velocity selection, a random sequence of the conditions was computer-generated for each participant. Participants walked on the treadmill, starting at 0.056 m/s, while velocity was increased in increments of 0.028 m/s at roughly 1 Hz. Participants were instructed to inform the examiner when the velocity of the treadmill was convenient and were allowed, if they so wished, to test velocities (increase or decrease velocity) before selection. This procedure was repeated for each condition according to the pre-generated random sequence. Once all velocities were obtained, testing commenced: a new random sequence was generated and participants walked, at their pre-selected velocity, in each condition according to this sequence. Between conditions the treadmill was stopped for 10 sec, during which, participants were instructed of their next condition. Each participant walked a total of 20 minutes.

Fixation targets for the DWG1, DWG3 and FG were marked as in the previous experiments. For DWGF, participants were instructed to fixate where they assumed their toes would be in the next step (i.e one step ahead). For the UR condition, participants were instructed to behave as they felt comfortable. Since the treadmill is elevated from the laboratory’s floor, a 3m×1.5m×0.2m wooden platform was custom built to create the illusion of continuity of the walking surface (with the targets for DWG1 and DWG3 attached to the platform). During all conditions, participants were instructed to try and walk in the middle of the treadmill to maintain a constant distance from the fixation targets. The Y coordinates of this position were obtained, beforehand, to control for any significant deviations during the experiment.

The treadmill used in this experiment was Mill (ForceLink, Culemborg, The Netherlands), a split-belt type treadmill with an embedded force-plate, able to record vertical ground reaction force (GRF) and COP trajectory. Data were collected continuously at 300Hz throughout the experiment. Since the treadmill is equipped with a front holding bar and its motors are mounted in the front, all experiments were executed in the treadmill’s reverse mode, thus, creating an imaginary, obstacle-free, walking path.

For processing, raw data from the treadmill’s force plate were exported to MATLAB. Using a dedicated script, individual time series was first smoothed with a moving average of 10 points, after which individual heel-strikes were identified: a surge in the vertical force, following the rapid change in COPX trajectory, exceeding body weight, was identified from the GRF and COPX time series. The heel-strike was defined as the time point at which maximal increase in force, along the surge, was identified. If the change in COPX was to the right, a right heel-strike followed and vise-versa. From this series of heel-strikes the parameter *step-time* was computed as the time elapsed between consecutive heel-strikes. The parameter *step-width* was also computed and defined as the distance, in mm, between the X coordinates of two consecutive heel-strikes.

Based on these computations we calculated center and spread measures for each trial and parameter, excluding the first and last 10 steps of each trial. Following the work of Chau et al. (2005) we evaluated the robustness of these measures by testing mean- and median-based measures. To make no assumptions we used the Wilcoxon signed-rank test. Results of these tests revealed significant differences (p<0.001) between mean and median values of all parameters and also between standard deviations (SD) and the median absolute deviations (MAD) of all parameters. Therefore, we decided to use the more robust estimates, i.e. median and MAD.

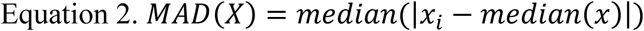

As primary outcome measures we computed the *finite-time* maximal Lyapunov exponent (i.e. divergence exponent, λ*) of the ground reaction force (Fz), COPY and COPX time series. Lyapunov exponents (λ) quantifies the sensitivity of a dynamical system to initial conditions. An n-dimensional dynamical system can be characterized by the rate at which close trajectories diverge over time. Under certain conditions, this rate of divergence along a given coordinate converges to an exponential relation, which can be simply described as follows:

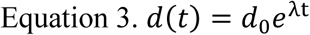

Where d represents the average separation as a function of time t, *d*_0_ is the initial separation and λ is the Lyapunov exponent of the system along this coordinate. Since the system is n-dimensional, it is characterized by an n-sized spectrum of lambdas. The maximal Lyapunov exponent (λ_max_) is simply the largest λ in the spectrum. As t grows larger, λ_max_ grows more and more dominant and the other exponents become negligible. When the evolution equation of the system is known, the Lyapunov spectrum is well defined in infinitesimal terms. However, estimating the Lyapunov spectrum of a hypothetical underlying system from a finite time series of measurements assumed to be produced thereby is by no means trivial. Various methods have been suggested for such estimation, but often it is only the maximal exponent that is of interest as a single indicator of the systems’ sensitivity to initial conditions. Rosenstein et al. (1993) proposed a relatively simple way of estimating just λ_max_ (denoted as λ*) from a finite time series. This method has been widely used in various fields of research, including human locomotion (Dingwell and Cusumano, 2000). Due to its simplicity, we adopted it too, although other methods (Wolf *et al.*, 1985) have been used as well.

To expose the Lyapunov spectrum and other dynamic system properties, a 1-dimensional series usually has to be “unfolded” or embedded in higher dimensions. This may be achieved by creating time-delayed copies of the original series (i.e. method of delays (Takens, 1981)). The data is then treated as multi-dimensional rather than 1-dimensional.

For walking, basically any time-dependent state variable/s may be used to characterize the dynamic properties of the system (i.e. the human body) using λ*. Any nearby positions of this variable, in state-space, represent nearly the same state and any one may be regarded as a perturbation of the other (Dingwell and Cusumano, 2000); thus, λ* may be used to quantify the rate at which these two positions diverge, representing the local dynamic steadiness (or more accurately, the unsteadiness) of gait.

To characterize the dynamics of walking in the present experiment, we computed λ* for the 5D space reconstructed from the Fz, COPY and COPX time series, using the middle 150 strides (Bruijn *et al.*, 2009). First, the individual unfiltered time series were time-normalized, using spline interpolation, to 200 points per stride so as to unify gait cycle and overall series lengths (Bruijn *et al.*, 2009). Next, the phase-space reconstruction interval was found by the mutual information first minimum method described by Fraser and Swinney (1986). Notably, values for the different state-variables (i.e. Fz, COPY and COPX) were very different. Using the “false nearest neighbors” (Kennel, Brown and Abarbanel, 1992), we tested a few examples and found that a minimal, and therefore optimal, dimension of 5 resulted for all. Others have found this number to be optimal for gait-derived data (Dingwell and Cusumano, 2000; England and Granata, 2007). We set the value of this parameter to 5 and refrained from computing it for each series separately.

Rosenstein’s average log divergence by lag graph was computed using the *Tisean* package (Hegger, Kantz and Schreiber, 1999), freely available at https://www.pks.mpg.de/~tisean/. Our MATLAB program only extracted the slope of the initial linear trend of this graph and plotted it. Since this slope is very loosely defined in the original paper, we tried various linear approximations. From these plots a prominent difference between the state-variables was observed: the initial rapid increase ends after roughly 1-1.5 steps, for the parameter Fz, after 1.5- 2 strides for the parameter COPX and even more for the parameter COPY. Moreover, significant fluctuations are easily observable throughout the plot. Therefore, approximation of the linear trend was greatly influenced by the time-period used. To overcome this difficulty, and for consistency, we did not constrain the linear approximation by time (i.e. to either step or stride), as is common in the literature (Bruijn *et al.*, 2013), but by magnitude. Namely, the approximation was computed over the 10%-90% range of the maximal value observed in the time series. The slope of this linear approximation (corresponding to the short-term λ* usually used in the literature) was used for statistical analysis.

Processing resulted in 150 data points for each parameter. These were exported to SPSS for statistical analysis. Since we anticipated a correlation between velocity and all other parameters, we first evaluated these correlations. Results of this procedure revealed correlations between step-time MAD, λ*COPY and λ*Fz, and velocity, but not for the other parameters (i.e. step-width MAD and λ*COPX). To control for the effect of velocity, we included walking velocity as a *fixed* effect in the statistical models of the appropriate parameters. Both the velocity uncontrolled and controlled models are reported. To test the effect of gaze position on walking steadiness we used mixed-effect models with subject as the *random* effect and condition as the *fixed* effect. The models’ residuals were then evaluated for normal distribution.

*Experiment 3-* 15 healthy young adults participated in this experiment. All experiments took place in a quiet, well-lit hallway, approximately 30 meters long and closed from both sides. The boundaries of the corridor (i.e. intersection of the walls and floor) were clearly marked using a patterned tape for later use. Participants were instructed to walk a 20-meter-long course, at a comfortable pace under two conditions: walking with their own shoes and walking with the Re- Step system (RS). The RS is elsewhere described (Bar-Haim *et al.*, 2013; Koren *et al.*, 2018; Koren, Parmet and Bar-Haim, 2019). Briefly, this is a shoe-like system with four pistons underneath each sole. In the passive mode these are aligned to create a plane parallel to the sole of the shoe. In the active mode, the pistons, during the swing phase, may change their length to create a plane oblique to the sole’s plane. These changes are perceived, during the stance phase, as walking on uneven surface. Naturally, these changes are visually unpredictable. To familiarize participants with the RS, they walked the entire course six times with the system in passive mode, before testing commenced.

For testing, participants performed 12 consecutive walks in each shoe type (i.e. two blocks consisting of 12 walks each). Before each walk, participants were presented with a tablet displaying either a simple countdown (from F to A) or a random sequence of digits, followed by a “GO” cue. The random sequence was displayed as a secondary, short-term memory task for purposes other than that of the current report. Therefore, only results for walks preceded by a simple countdown are reported here (i.e. 6 walks in each condition). Gaze positions were monitored throughout the experiment, using binocular eye-tracking glasses (ETG, SensoMotoric Instruments, Teltow, Germany), at a 60 Hz sampling rate. Before testing, the ETG was calibrated according to the manufacturer’s instructions, using a 3-point calibrating procedure. Using the software provided by the manufacturer (BeGaze, v.3.5), gaze positions were superimposed on a scenery video, from a front-mounted camera. Each video was then manually screened and calibration of the ETG, before each walk, was evaluated. If needed, post-hoc calibration was performed assuming participants fixated on the middle of the tablet. Following this procedure, the scenery video and gaze positions data were exported for further analysis.

Following the procedure described above, the recorded video and raw gaze data were processed using MATLAB. Before processing the data, we initially extracted two image patches at locations of the tapes that marked the corridor boundaries. The extraction was done manually by selecting a line segment over each marker tape in a single frame and extracting the patches at the middle point of each line segment. With the two patches at hand, the video was processed automatically in a frame-by-frame manner.

In each frame, we used the manually extracted patches as templates to detect regions that contained the striped pattern of the marker tapes. To this end, we used a template-matching technique based on normalized cross-correlation in the frequency domain between the patches and the frame. This process resulted in a grayscale map with higher values indicating regions that more likely belong to the marker tapes. By thresholding this map, we obtained a binary image indicating regions in the marker tapes.

Finally, we used the Hough transform to find the parameters of a straight line passing through these regions. Once the marker tapes were detected and represented as straight lines in the frame, we estimated the gaze distance based on the known corridor width (i.e., the distance between the marker tapes) and using perspective geometry calculations. Further, from these calculations the intersection point of the lines was determined and defined as the “vanishing point”. Raw gaze coordinates were then referenced to this point and the vertical distance and its variability were calculated. These values are presented as percentage of the frame size. For simplicity, we used the inverse of the vertical gaze distance values such that large values indicate a gaze directed straight ahead (i.e. toward the horizon) and small values indicate DWG.

Since gaze-position data is highly variable, we used median and median-absolute-deviation (MAD), as center and spread measures, respectively. As our main outcome measure we used gaze position distance from the vanishing point, on the vertical axis (vertical gaze position), and as a secondary measure we used gaze distance. Although both of these outcome measures quantify the same behavior (but in different scales), the former may be computed for every sample while the latter is meaningful (in the context of stepping control) only when gaze was directed onto the future path. Other outcome measures included the look-ahead distance variability, walking velocity and the proportion of gaze positions (from the total number of samples) directed onto the future path.

For statistical analysis we used a linear mixed-effect model with participants as the *random* effect and condition as the *fixed* effect. However, given that the distributions of the two variables, gaze distance and its variability, were skewed to the right, analysis was performed using Gamma distribution instead of normal distribution. In all cases the models’ residuals were evaluated for normal distribution.

## Acknowledgments

This research was supported by the Helmsley Charitable Trust through the Agricultural, Biological and Cognitive Robotics Initiative and by the Marcus Endowment Fund both at Ben-Gurion University of the Negev.

## Bibliography

Aoki, O., Otani, Y., Morishita, S. and Domen, K. (2017) ‘The effects of various visual conditions on trunk control during ambulation in chronic post stroke patients’, Gait & posture, 52, pp. 301-307. doi: S0966-6362(16)30706-8 [pii].

Aoki, O., Otani, Y., Morishita, S. and Domen, K. (2014) ‘Influence of gaze distance and downward gazing on postural sway in hemiplegic stroke patients’, Experimental brain research, 232(2), pp. 535-543. doi: 10.1007/s00221-013-3762-3 [doi].

Bardy, B.G., Warren, W.H.,Jr and Kay, B.A. (1996) ‘Motion parallax is used to control postural sway during walking’, Experimental brain research, 111(2), pp. 271-282. doi: 10.1007/bf00227304 [doi].

Bar-Haim, S., Harries, N., Hutzler, Y., Belokopytov, M. and Dobrov, I. (2013) ‘Training to walk amid uncertainty with Re-Step: measurements and changes with perturbation training for hemiparesis and cerebral palsy’, Disability and rehabilitation.Assistive technology, 8(5), pp. 417-425. doi: 10.3109/17483107.2012.754954 [doi].

Bruijn, S.M., Meijer, O.G., Beek, P.J. and van Dieen, J.H. (2013) ‘Assessing the stability of human locomotion: a review of current measures’, Journal of the Royal Society, Interface, 10(83), pp. 20120999. doi: 10.1098/rsif.2012.0999 [doi].

Bruijn, S.M., van Dieen, J.H., Meijer, O.G. and Beek, P.J. (2009) ‘Statistical precision and sensitivity of measures of dynamic gait stability’, Journal of neuroscience methods, 178(2), pp. 327-333. doi: 10.1016/j.jneumeth.2008.12.015 [doi].

Buckley, J.G., Anand, V., Scally, A. and Elliott, D.B. (2005) ‘Does head extension and flexion increase postural instability in elderly subjects when visual information is kept constant?’, Gait & posture, 21(1), pp. 59-64. doi: S0966636203002017 [pii].

Chau, T., Young, S. and Redekop, S. (2005) ‘Managing variability in the summary and comparison of gait data’, Journal of neuroengineering and rehabilitation, 2, pp. 22-0003-2-22. doi: 1743-0003-2-22 [pii].

Clark, J.E. (1995) ‘On becoming skillful: patterns and constraints’, Research quarterly for exercise and sport, 66(3), pp. 173-183. doi: 10.1080/02701367.1995.10608831 [doi].

Collins, J.J. and De Luca, C.J. (1993) ‘Open-loop and closed-loop control of posture: a random-walk analysis of center-of-pressure trajectories’, Experimental brain research, 95(2), pp. 308–318.

Collins, J.J., De Luca, C.J., Burrows, A. and Lipsitz, L.A. (1995) ‘Age-related changes in open-loop and closed-loop postural control mechanisms’, Experimental brain research, 104(3), pp. 480-492. doi: 10.1007/bf00231982 [doi].

Dingwell, J.B. and Cusumano, J.P. (2000) ‘Nonlinear time series analysis of normal and pathological human walking’, Chaos (Woodbury, N.Y.), 10(4), pp. 848-863. doi: 10.1063/1.1324008 [doi].

Domínguez-Zamora, F.J., Gunn, S.M. and Marigold, D.S. (2018) ‘Adaptive Gaze Strategies to Reduce Environmental Uncertainty During a Sequential Visuomotor Behaviour’, Scientific reports, 8(1), pp. 14112.

Ellmers, T.J., Cocks, A.J. and Young, W.R. (2019) ‘Evidence of a link between fall-related anxiety and high-risk patterns of visual search in older adults during adaptive locomotion’, The journals of gerontology.Series A, Biological sciences and medical sciences,. doi: glz176 [pii].

Ellmers, T.J. and Young, W.R. (2019) ‘The influence of anxiety and attentional focus on visual search during adaptive gait’, Journal of experimental psychology.Human perception and performance, 45(6), pp. 697-714. doi: 10.1037/xhp0000615 [doi].

England, S.A. and Granata, K.P. (2007) ‘The influence of gait speed on local dynamic stability of walking’, Gait & posture, 25(2), pp. 172-178. doi: S0966-6362(06)00041-5 [pii].

Fraser, A.M. and Swinney, H.L. (1986) ‘Independent coordinates for strange attractors from mutual information’, Physical review.A, General physics, 33(2), pp. 1134-1140. doi: 10.1103/physreva.33.1134 [doi].

Guerraz, M. and Bronstein, A.M. (2008) ‘Ocular versus extraocular control of posture and equilibrium’, Neurophysiologie clinique = Clinical neurophysiology, 38(6), pp. 391-398. doi: 10.1016/j.neucli.2008.09.007 [doi].

Guerraz, M., Sakellari, V., Burchill, P. and Bronstein, A.M. (2000) ‘Influence of motion parallax in the control of spontaneous body sway’, Experimental brain research, 131(2), pp. 244-252. doi: 10.1007/s002219900307 [doi].

Hayhoe, M.M. and Matthis, J.S. (2018) ‘Control of gaze in natural environments: effects of rewards and costs, uncertainty and memory in target selection’, Interface focus, 8(4), pp. 20180009.

Hegger, R., Kantz, H. and Schreiber, T. (1999) ‘Practical implementation of nonlinear time series methods: The TISEAN package’, Chaos (Woodbury, N.Y.), 9(2), pp. 413-435. doi: 10.1063/1.166424 [doi].

Hollands, M.A., Marple-Horvat, D.E., Henkes, S. and Rowan, A.K. (1995) ‘Human Eye Movements During Visually Guided Stepping’, Journal of motor behavior, 27(2), pp. 155-163. doi: 10.1080/00222895.1995.9941707 [doi].

Kapoula, Z. and Le, T.T. (2006) ‘Effects of distance and gaze position on postural stability in young and old subjects’, Experimental brain research, 173(3), pp. 438-445. doi: 10.1007/s00221-006-0382-1 [doi].

Kay, B.A. and Warren Jr, W.H. (2001) ‘Coupling of posture and gait: mode locking and parametric excitation’, Biological cybernetics, 85(2), pp. 89–106.

Kennel, M.B., Brown, R. and Abarbanel, H.D. (1992) ‘Determining embedding dimension for phase-space reconstruction using a geometrical construction’, Physical review.A, Atomic, molecular, and optical physics, 45(6), pp. 3403-3411. doi: 10.1103/physreva.45.3403 [doi].

Koren, Y., Raanan, Y., Parmet, Y. and Bar-Haim, S. (2018) ‘Treading on the unknown-the feasibility of a novel approach to investigating the motor control of walking’, Physiological Measurement, 39(4), pp. 04NT01-6579/aab659. doi: 10.1088/1361-6579/aab659 [doi].

Koren, Y., Parmet, Y. and Bar-Haim, S. (2019) ‘Treading on the unknown increases prefrontal activity: A pilot fNIRS study’, Gait & Posture, 69, pp. 96-100. doi: https://doi.org/10.1016/j.gaitpost.2019.01.026.

Lee, D.N. and Lishman, J. (1975) ‘Visual proprioceptive control of stance.’, Journal of human movement studies,.

Marigold, D.S. and Patla, A.E. (2007) ‘Gaze fixation patterns for negotiating complex ground terrain’, Neuroscience, 144(1), pp. 302–313.

Marigold, D.S. (2008) ‘Role of peripheral visual cues in online visual guidance of locomotion’, Exercise and sport sciences reviews, 36(3), pp. 145-151. doi: 10.1097/JES.0b013e31817bff72 [doi].

Matthis, J.S. and Fajen, B.R. (2013) ‘Humans exploit the biomechanics of bipedal gait during visually guided walking over complex terrain’, Proceedings of the Royal Society B: Biological Sciences, 280(1762), pp. 20130700.

Matthis, J.S., Barton, S.L. and Fajen, B.R. (2015) ‘The biomechanics of walking shape the use of visual information during locomotion over complex terrain’, Journal of vision, 15(3), pp. 10.1167/15.3.10. doi: 10.1167/15.3.10 [doi].

Matthis, J.S. and Fajen, B.R. (2014) ‘Visual control of foot placement when walking over complex terrain’, Journal of experimental psychology.Human perception and performance, 40(1), pp. 106-115. doi: 10.1037/a0033101 [doi].

Matthis, J.S., Yates, J.L. and Hayhoe, M.M. (2018) ‘Gaze and the Control of Foot Placement When Walking in Natural Terrain’, Current biology : CB, 28(8), pp. 1224-1233.e5. doi: S0960-9822(18)30309-9 [pii].

Melzer, I., Benjuya, N. and Kaplanski, J. (2004) ‘Postural stability in the elderly: a comparison between fallers and non-fallers’, Age and Ageing, 33(6), pp. 602-607. doi: 33/6/602 [pii].

Melzer, I., Kurz, I. and Oddsson, L.I. (2010) ‘A retrospective analysis of balance control parameters in elderly fallers and non-fallers’, Clinical biomechanics (Bristol, Avon), 25(10), pp. 984-988. doi: 10.1016/j.clinbiomech.2010.07.007 [doi].

Patla, A.E. and Vickers, J.N. (2003) ‘How far ahead do we look when required to step on specific locations in the travel path during locomotion?’, Experimental brain research, 148(1), pp. 133-138. doi: 10.1007/s00221-002-1246-y [doi].

Patla, A.E. and Vickers, J.N. (1997) ‘Where and when do we look as we approach and step over an obstacle in the travel path?’, Neuroreport, 8(17), pp. 3661-3665. doi: 10.1097/00001756-199712010-00002 [doi].

Patla, A.E. (1997) ‘Understanding the roles of vision in the control of human locomotion’, Gait & posture, 5(1), pp. 54–69.

Peterka, R.J. (2000) ‘Postural control model interpretation of stabilogram diffusion analysis’, Biological cybernetics, 82(4), pp. 335-343. doi: 10.1007/s004220050587 [doi].

Prieto, T.E., Myklebust, J.B., Hoffmann, R.G., Lovett, E.G. and Myklebust, B.M. (1996) ‘Measures of postural steadiness: differences between healthy young and elderly adults’, IEEE transactions on bio-medical engineering, 43(9), pp. 956-966. doi: 10.1109/10.532130 [doi].

Reynolds, R.F. and Day, B.L. (2005a) ‘Visual guidance of the human foot during a step’, The Journal of physiology, 569(2), pp. 677–684.

Reynolds, R. and Day, B. (2005b) ‘Rapid visuo-motor processes drive the leg regardless of balance constraints’, Current Biology, 15(2), pp. R48–R49.

Rosenstein, M.T., Collins, J.J. and De Luca, C.J. (1993) ‘A practical method for calculating largest Lyapunov exponents from small data sets’, Physica D: Nonlinear Phenomena, 65(1-2), pp. 117–134.

Salinas, M.M., Wilken, J.M. and Dingwell, J.B. (2017) ‘How humans use visual optic flow to regulate stepping during walking’, Gait & posture, 57, pp. 15-20. doi: S0966-6362(17)30184-4 [pii].

Schrater, P.R., Knill, D.C. and Simoncelli, E.P. (2001) ‘Perceiving visual expansion without optic flow’, Nature, 410(6830), pp. 816–819.

Smid, K. and Den Otter, A. (2013) ‘Why you need to look where you step for precise foot placement: the effects of gaze eccentricity on stepping errors’, Gait & posture, 38(2), pp. 242–246.

Stoffregen, T.A. (1985) ‘Flow structure versus retinal location in the optical control of stance’, Journal of experimental psychology. Human perception and performance, 11(5), pp. 554-565. doi: 10.1037//0096-1523.11.5.554 [doi].

Takens, F. (1981) ‘Detecting strange attractors in turbulence’Dynamical systems and turbulence, Warwick 1980 Springer, pp. 366–381.

Ustinova, K. and Perkins, J. (2011) ‘Gaze and viewing angle influence visual stabilization of upright posture’, Brain and behavior, 1(1), pp. 19-25. doi: 10.1002/brb3.10 [doi].

Warren Jr, W.H. (1995) ‘Self-motion: Visual perception and visual control’Perception of space and motion Elsevier, pp. 263–325.

Warren, W.H., Jr, Kay, B.A., Zosh, W.D., Duchon, A.P. and Sahuc, S. (2001) ‘Optic flow is used to control human walking’, Nature neuroscience, 4(2), pp. 213-216. doi: 10.1038/84054 [doi].

Warren, W.H., Kay, B.A. and Yilmaz, E.H. (1996) ‘Visual control of posture during walking: functional specificity’, Journal of experimental psychology. Human perception and performance, 22(4), pp. 818-838. doi: 10.1037//0096-1523.22.4.818 [doi].

Wolf, A., Swift, J.B., Swinney, H.L. and Vastano, J.A. (1985) ‘Determining Lyapunov exponents from a time series’, Physica D: Nonlinear Phenomena, 16(3), pp. 285–317.

Wolsley, C., Sakellari, V. and Bronstein, A. (1996) ‘Reorientation of visually evoked postural responses by different eye-in-orbit and head-on-trunk angular positions’, Experimental brain research, 111(2), pp. 283–288.

Zietz, D. and Hollands, M. (2009) ‘Gaze behavior of young and older adults during stair walking’, Journal of motor behavior, 41(4), pp. 357-365. doi: 10.3200/JMBR.41.4.357-366 [doi].

